# Parsing the effect of co-culture with brain organoids on Diffuse Intrinsic Pontine Glioma (DIPG) using quantitative proteomics

**DOI:** 10.1101/2023.05.19.541551

**Authors:** Victoria G Prior, Simon Maksour, Sara Miellet, Amy J Hulme, Yuyan Chen, Mehdi Mirzaei, Yunqi Wu, Mirella Dottori, Geraldine M O’Neill

## Abstract

Diffuse Intrinsic Pontine Gliomas (DIPGs) are deadly brain cancers in children for which there is currently no effective treatment. This can partly be attributed to preclinical models that lack essential elements of the *in vivo* tissue environment, resulting in treatments that appear promising preclinically, but fail to result in effective cures. Recently developed co-culture models combining stem cell-derived brain organoids with brain cancer cells provide tissue dimensionality and a human-relevant tissue-like microenvironment. As these models are technically challenging and time consuming it is imperative to establish whether interaction with the organoid influences DIPG biology and thus warrants their use. To address this question, we cultured DIPG cells with cortical organoids. We created “mosaic” co-cultures enriched for tumour cell-neuronal cell interactions versus “assembloid” co-cultures enriched for tumour cell-tumour cell interactions. Sequential window acquisition of all theoretical mass spectra (SWATH-MS) was used to analyse the proteomes of DIPG fractions isolated by flow-assisted cell sorting. Control proteomes from DIPG spheroids were compared with DIPG cells isolated from mosaic and assembloid co-cultures. This revealed that tumour cell adhesion was reduced, and DNA synthesis and replication were increased, in DIPG cells under either co-culture condition. By contrast, the mosaic co-culture was associated with pathways implicated in dendrite growth. We propose that co-culture with brain organoids is a valuable tool to parse the contribution of the brain microenvironment to DIPG tumour biology.

## INTRODUCTION

DIPG is a devastating paediatric brain tumour for which no effective treatments exist. DIPG tumours rarely, if ever, metastasise outside of the brain, but aggressively and diffusely invade neighbouring healthy tissue, displacing, distorting and destroying white matter tracts ^1^. Patient survival rates have not improved over the past 40 years ^2^, and no chemotherapeutic agents have proven to be effective ^3^. Fractionated radiotherapy remains the mainstay treatment, but provides only transient relief of symptoms and no significant contribution to overall- or progression-free survival ^4^. More recently, the development of patient-derived DIPG lines from autopsy and biopsy samples ^5^ has started to yield important new insights into the genetic landscape of this tumour. Despite recent highlights underscoring the importance of the brain tissue context to DIPG specifically ^6^ and more broadly to high grade gliomas ^7–9^, understanding of how DIPG interaction with normal brain cells impacts DIPG biology is still limited, but is likely a key factor in the successful eradication of DIPG tumours. The incorporation of normal brain tissue into preclinical models of DIPG may thus improve preclinical investigations, leading to improved patient outcomes.

The highly invasive nature of DIPG impedes successful treatment. Similarly, glioblastomas (GBM) disseminate widely throughout the healthy brain tissue with the migratory GBM cells having stem-like features and being associated with tumour recurrence ^10–12^. As cell invasion and migration is determined by the biochemical and structural features of the surrounding tissue, the inclusion of a brain tissue-like environment into preclinical models should encourage DIPG dissemination. Co-culture of patient tumour cells with human pluripotent stem cell (PSC)-derived brain organoids offers an *in vitro* platform for modelling multiple aspects of *in vivo* tumour biology, most importantly, invasion and dissemination. GBM cells spontaneously invade into brain organoids ^13–15^, are more resistant to typically used GBM therapies, versus the same GBM cells grown in 2-dimensional (2D) culture^13^ and retain the cellular heterogeneity of the primary tumours^16^. Therefore, we reasoned that brain organoids would likely provide a platform for investigating invasive DIPG.

The preparation of PSC-derived brain organoids is technically challenging and, depending on the desired level of maturity and fate, can take many months. To determine the value of this model for advancing the understanding of DIPG we investigated whether co-culture with brain organoids influences DIPG biology. The main tumour bulk in which tumour cell-tumour cell interactions predominate was analysed by combining pre-formed DIPG spheroids with cortical organoids (assembloids). The diffusely invaded DIPG in which tumour cell-tumour microenvironment (TME) interactions predominate was achieved by mixing and then reforming dissociated spheroids and cortical organoids (mosaics). Proteomes of DIPG tumour fractions from each model were collected by sequential window acquisition of all theoretical fragment ion spectra mass spectrometry (SWATH-MS) and compared with proteomes of DIPG spheroids. The data revealed that the organoids support DIPG invasion and influence cellular signalling programs.

## RESULTS

### DIPG co-cultures with cortical organoids

To analyse the effect of a healthy brain microenvironment on DIPG cells we used two approaches to co-culturing DIPG with dorsal-cortically-fated brain organoids derived from human embryonic stem cells (hESC) (see schematic, Figure 1A). One approach involved formation of assembloids consisting of pre-formed DIPG spheroids combined with cortical organoids, designed to mimic the primary tumour bulk, with the interface between the DIPG spheroid and the cortical organoid representing the invasive front of the tumour. In the second approach, mosaics were formed by dissociating individual pre-formed DIPG spheroids and organoids, then mixing the cells together and allowing the structures to reform, similar to previous reports for GBM^17^. The mosaic model represents cells that have diffusely invaded the healthy brain tissue and this approach was also taken to provide sufficient material for protein extraction and proteomic analyses. In each model, hESC were subject to a differentiation protocol for 16 days (Figure 1B), at which point the pluripotency marker OCT4 was lost and the neural progenitor SOX2 was upregulated (Figure 1C). Concurrently, DIPG24 spheroids were formed for the last 5 days of the organoid maturation protocol (Figure 1D). Assembloids were created by combining organoids and tumour spheroids in a 1:1 ratio and mosaic cultures were created mixing equal numbers of dissociated organoid and spheroid cells (Figure 1E).

**Figure 1.**
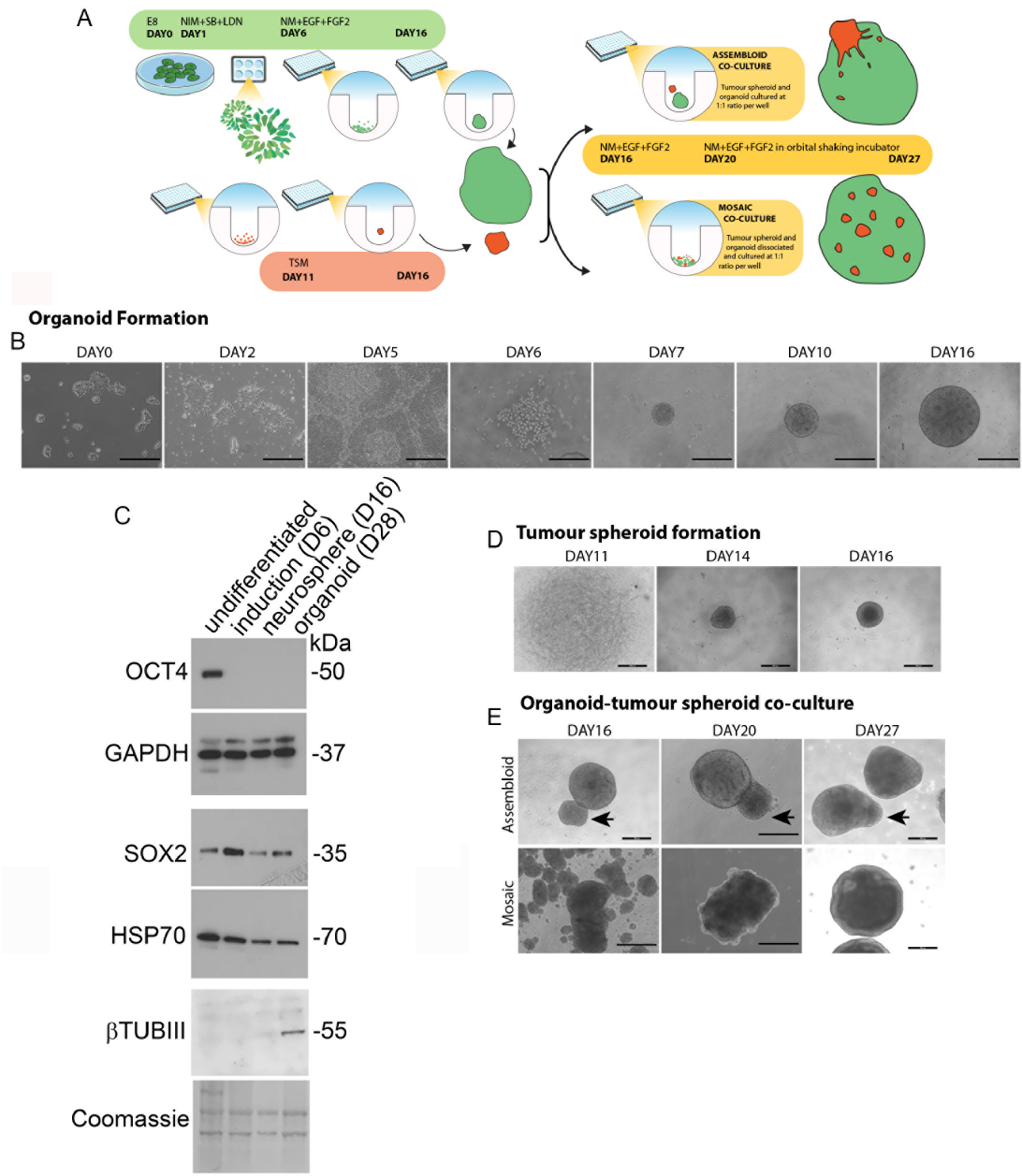
Formation of assembloid and mosaic co-cultures. A. Schematic overview of co-culture establishment. Assembloids were formed by combining day 16 cortical organoids with pre-formed HGG spheroids in a 1:1 ratio for 3 days, followed by further culture in a shaking incubator up to day 27. Mosaics are formed by combining dissociated day 16 cortical organoids with dissociated HGG spheroids (1spheroid:1organoid) for 3 days, then transfer to shaking incubator up to day 27. B. Bright field images showing typical formation of organoids showing the progression of H9 human embryonic stem cells (hESCs) to neural rosette formation (day 0-6), followed by harvesting and replating to induce neurospheres (day 7-16). C. Western blots of protein extracts from H9 cells under the indicated conditions: undifferentiated = H9 stem cells; induction = Day 6; neurosphere = Day 16; organoid = Day 28. Lysates were probed for expression of the pluripotency markers OCT4, and SOX2 and βTUBIII. GAPDH, HSP70 and HSP90 = loading controls. D. HGG cells (plated on day 11 of the cortical induction protocol) compacted over 5 days to form tumour spheroids. E. At Day 16 of the organoid differentiation protocol cultures were combined as outlined in panel A to form assembloids or mosaics. Arrows indicate DIPG spheroid position in the assembloid. Scale bar = 500µm.

Importantly, the reformed organoids in the mosaic model retained the expected morphological and cellular features. Comparison of control organoids with dissociated/re-associated organoids confirmed similar expression and spatial distribution of vimentin, SOX2, PAX6, TBR2, βTUBIII and MAP2 (Figure 2A). Neural rosettes were apparent in the reformed organoids, although they were more numerous and smaller on average than the control organoid counterparts (Figure 2B). The mosaic organoids therefore retained the key features of cortically fated organoids.

**Figure 2.**
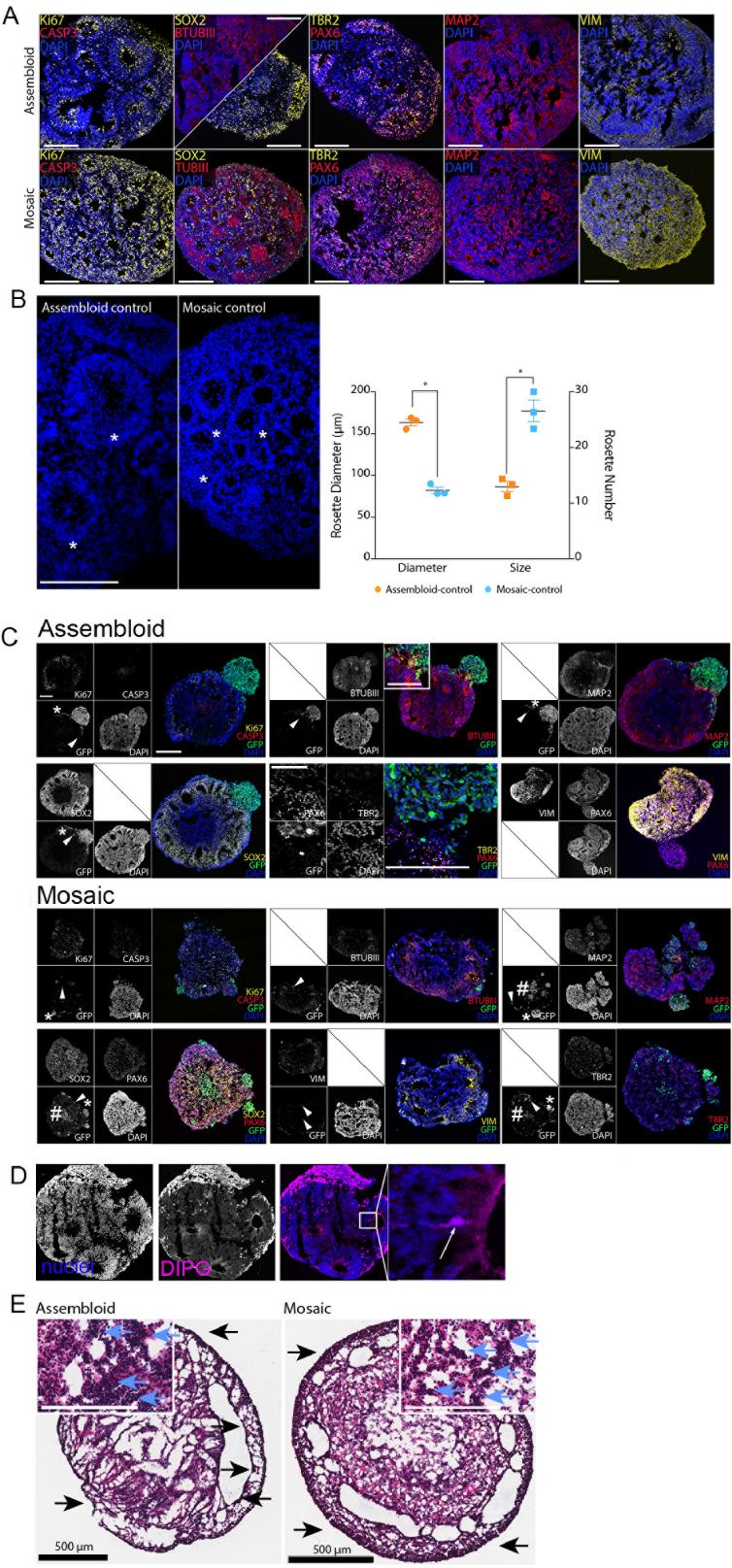
Organoid structure and molecular composition is maintained in co-cultures. A. Maximum z-projections of confocal slices show similar expression and distribution of neural stem cell markers vimentin (VIM, yellow) and SOX2 (yellow), ventricular zone progenitor PAX6 (red) and subventricular zone progenitor TBR2 (yellow) in control organoids (assembloid) and disaggregated/reaggregated organoids (mosaic). Markers are collectively indicative of forebrain identity. Beta-tubulin III (TUBIII, red) and MAP2 (red) highlight neuronal structures. Images shown are representative examples from n=3 independent repeats. Scale bars = 200 µm. B. Hoescht-blue stained images highlighting characteristic neural rosettes in control organoids (assembloid control) and disaggregated/reaggregated organoids (mosaic control). Asterisks mark the centre of example neural rosettes. Graph shows the average rosette diameter and number for the indicated conditions. Data points show the means ± SEM from 3 independent replicates, n = 3 organoids per replicate. \**P* = 0.0002 (diameter) and \**P* = 0.0031 (number) Students’ *t*-test. C. H9-derived organoids co-cultured with DIPG24 cells transduced to express eGFP. In assembloid cultures, arrow heads indicate GFP-positive tumour cells that have invaded into the body of the organoid and asterisks highlight examples of GFP-positive DIPG24 cells that have invaded along the organoid boundary. In mosaic cultures, asterisks highlight small, rounded clusters of GFP-positive DIPG24 cells, arrow heads indicate individual cells and hash tags indicate loose cell clusters. Maximum z-projections of confocal z-slices immunostained for dorsal cortical markers as indicated. Sections additionally immunostained for proliferation (Ki67, yellow) and apoptosis (cleaved caspase 3, CASP3, red). Images shown are representative examples from n = 3 independent replicates, Scale bar = 200 µm. D. Assembloid showing nuclei (blue) and GFP-positive DIPG cells (pink). Panel on the right-hand side shows a magnified image highlighting an example of a DIPG cell incorporated into a neural rosette (arrow). E. Hematoxylin and eosin-stained sections 17 days post-co-culture. Arrows show basophilic staining of tumours cells that have extensively perfused the organoid (blue arrows, insets) and tumour cells encapsulating the periphery of the organoids (black arrows). Scale bars = 500 µm, insets = 200 µm.

In assembloids, DIPG24 cells disseminated away from the spheroid and invaded throughout the organoid (Figure 2C, assembloids, arrowhead) and along the organoid surface (Figure 2C, assembloids, asterisk). In some instances, cells integrated into the neural rosettes (Figure 2D, inset). In mosaics, GFP-positive cells were distributed either as small, rounded clusters (Figure 2C, mosaics, asterisks), loose dispersed aggregates (Figure 2C, mosaics, hash tags) or single cells (Figure 2C, mosaics, arrow heads). Immunostaining for SOX2, ΒTUBIII, PAX6, MAP2 and VIM confirmed that the organoids retained the key structural and molecular features in the co-cultures (Figure 2C). The cortical organoids therefore provide an organised, permissive tissue environment that supports physiologically relevant DIPG invasion. Conversely, U87MG GBM cells, that do not invade health brain tissue in orthotopic mouse models ^18, 19^, neither invaded brain organoids in the assembloid model, nor incorporated into organoids in the mosaic model (Supplementary Figure 1). Notably, longer periods of DIPG/organoid co-culture resulted in the tumour cells overtaking the organoids in either co-culture model (Figure 2E), therefore the time of co-culture was limited to 10 days.

### Decreased adhesion and increased DNA synthesis and replication in DIPG co-cultured with cortical organoids

To determine the influence of the cortical organoid microenvironment on signalling we analysed DIPG cellular proteomes by SWATH-MS of proteins extracted from the co-cultures. Purified DIPG24 cell fractions from assembloids and mosaics were collected by disassociation and flow-assisted cell sorting into GFP+ve (DIPG24, >70% and >89% purity from the assembloid and mosaic cultures, respectively, Supplementary Figure 2) and GFP-ve (H9-derived organoid) (>99% purity from either co-culture model, Supplementary Figure 2) populations. In parallel, control protein extracts were independently prepared from DIPG24 spheroids, organoids (assembloid organoid control) and disassociated/reassociated organoids (mosaic organoid control). Details of samples and protein quantification are shown in Supplementary Figure 3. Following SWATH-MS, protein expression and abundance was calculated to be the same between replicates in control cultures (cortical organoids, disassociated/reassociated organoid and spheroids) validating the SWATH-MS sample preparation (Figure 3A).

**Figure 3.**
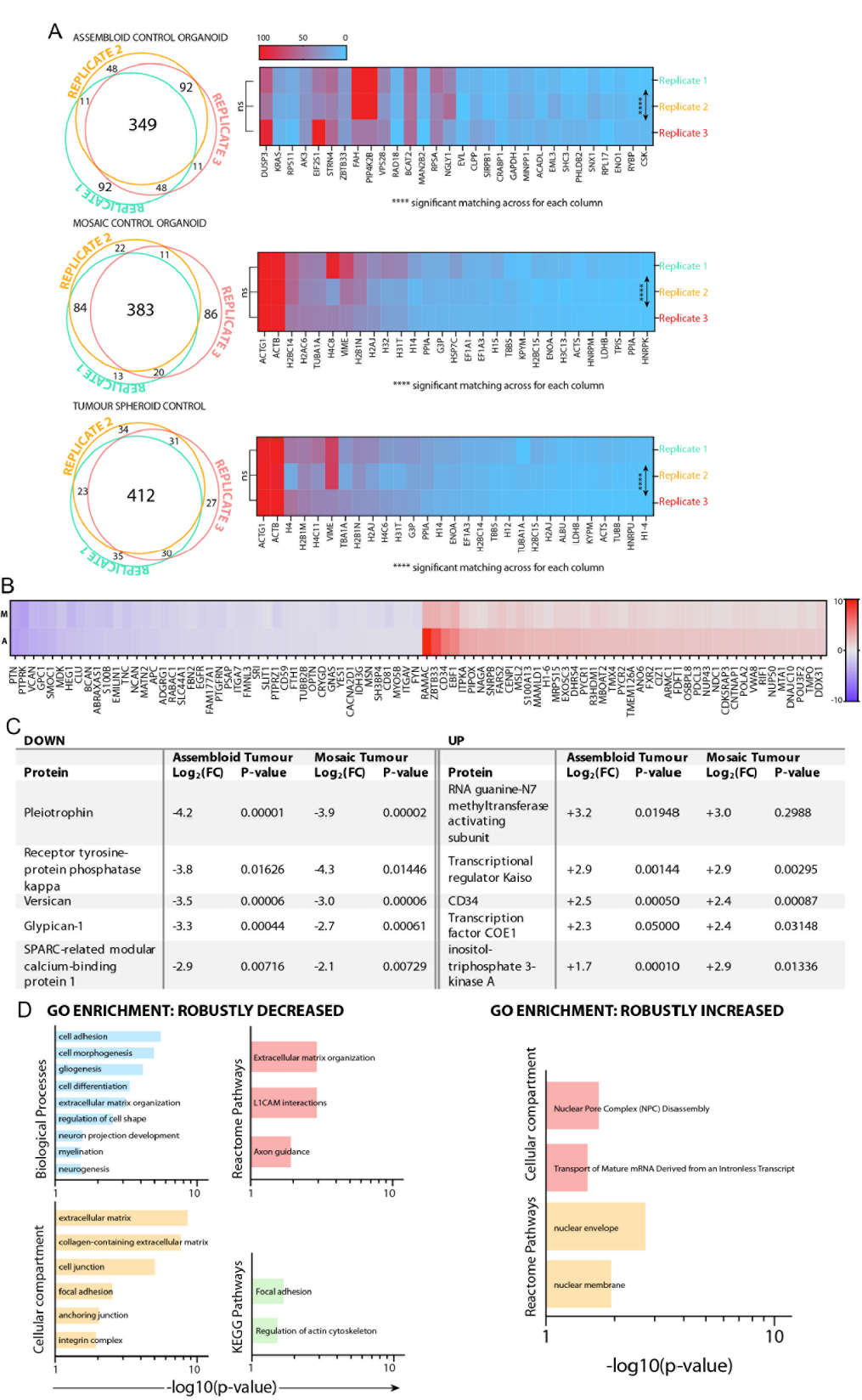
Co-culture with cortical organoids affects cancer-relevant signalling in DIPG24 cells. A. Spectral ion libraries were compiled from all samples. The abundance of peptides meeting quality control and SWATH-MS conditions was quantitated for all conditions. Shown are comparisons for peptides purified from cortical organoids (assembloid control organoids), disassociated/reassociated organoids (mosaic control organoids) and DIP24 spheroids (tumour spheroid controls) from 3 independent biological repeats. Venn diagrams show the overlap of protein expression from the top 500 most abundant proteins identified in each replicate. Heat maps show the normalized abundance of the top 30 identified proteins for each replicate, where red indicates the highest expression and blue indicates the lowest. There was no significant difference between replicates, while there was significant matching (****p<0.0001) across columns within each condition, indicating similar protein abundance across replicates. One-way ANOVA within subjects with Geisser-Greenhouse correction for matched data, n = 30 analysed per replicate. B. Heat map comparing the fold change (log10 [fold change]) for DIPG24 proteins that were altered in both assembloid and mosaic cultures. M = mosaic co-culture and A = assembloid co-culture. C. Proteins displaying the greatest fold change (log2 [fold change]) that were common to DIP24 cells from either assembloid or mosaic cultures. D. Gene ontology (GO) enrichment analyses of proteins that were commonly decreased (43 proteins) or increased (44 proteins). The respective GO lists were clustered by biological process (blue), KEGG pathways (green), cellular compartment (orange) and reactome (red) pathways. Bar charts show the significant (-log10 [p-value]) for each term.

In total, 87 proteins were commonly regulated (43 downregulated and 44 upregulated) in DIPG cells from either co-culture model, when compared with the proteomes of control DIPG spheroids (Figure 3B). Details of the top 5 proteins with the greatest fold decrease and increase are shown (Figure 3C). Gene ontology (GO) analyses were performed using the list of common DIPG24 expression changes. Protein expression terms associated with nuclear pore complex disassembly, increased transport of intron-less mature mRNA transcript were increased, while terms associated with cell adhesion were decreased (Figure 3D). The observation that there are pathways commonly altered under either co-culture condition suggests a coordinated program of signalling that is stimulated by the organoids, that does not require direct contact between DIPG and organoid cells.

The putative decrease in cell adhesion stimulated by the cortical organoid microenvironment suggested by the proteomic analyses was further analysed. Assembloids were immunostained for the adhesion marker β1 integrin distribution. This sub-unit forms heterodimers with a wide variety of α-subunits and thus provides a readout for multiple heterodimers. In control spheroids, β1 integrin is enriched at the spheroid periphery (Figure 4A). By contrast, there is little evidence of β1 integrin enrichment at the spheroid periphery in assembloids. Moreover, DIPG24 cells that have invaded into the organoid have lost membrane-associated β1 integrin expression (Figure 4A). These data thus support the proteomic pathway analyses suggesting that co-culture with cortical organoids decreases DIPG cell adhesion.

**Figure 4.**
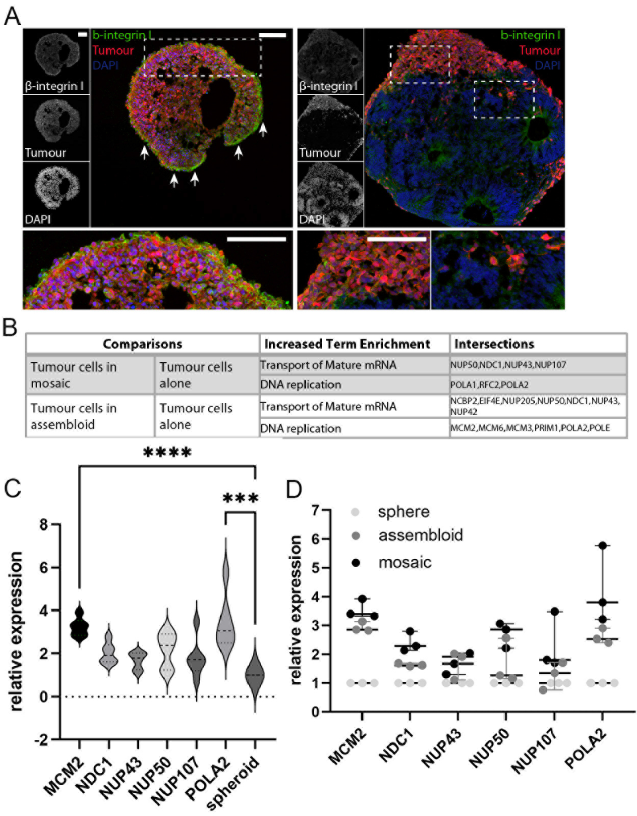
Co-culture with organoids reduces adhesion and increases MCM2 and POLA2 expression. A. Maximum intensity z-projections of confocal sections of DIPG24 spheroids (left panels) and DIPG24/cortical organoid assembloids (right panels) immunostained with anti-β1 integrin antibodies (green). GFP-positive tumour cells are false-coloured red. Dotted boxed regions shown magnified below each image. Arrows indicate cells on the periphery with highly organized β1-integrin at the cell boundaries. Images shown are representative of 3 independent repeats. Scale bars = 100 µm. B. Table summarizing GO term enrichments and gene product intersections for each term. Terms are described for group (left) relative to comparison group (right). C. RT-qPCR analysis of the indicated gene products in DIPG24 cells isolated from assembloids and mosaics. n = 3 biological replicates, one-way ANOVA performed relative to DIPG spheroid condition. \*\*\*\**P*<0.0001, *** *P*<0.001. D. Expression data from (C), showing individual data points from DIPG24 cells grown in mosaic or assembloid conditions, as indicated.

To further confirm the signalling program stimulated by organoid co-culture, representative GO gene intersections were selected for quantification (Figure 4B): minichromosome maintenance complex component 2 (*MCM2*); nucleoporin *NDC1*; nucleoporin 43 (*NUP43*); *NUP50*; *NUP107*; and DNA polymerase alpha 2 (*POLA2*). RNA quantification revealed that organoid co-culture induced *MCM2* and *POLA2* in DIPG24 cells (Figure 4C). Expression of each of the 6 genes tended to be highest in the mosaics, although the difference was not significant (Figure 4D). Collectively, the data suggest that the cortical organoid microenvironment results in decreased membrane localisation of integrins and increased expression of *MCM2* and *POLA2* in DIPG24 cells.

### Changes unique to DIPG cells in assembloids versus mosaics

To identify potential changes associated with contact between DIPG cells and the cortical organoid cells, we assessed unique protein changes between the two models. Proteomes were first filtered to identify proteins that were significantly altered in the assembloid or mosaic cultures, respectively, versus spheroid cultures alone (Figure 5A). From this list of proteins, proteins that were significantly changed in one co-culture condition, while being unchanged in the other co-culture condition were determined. The top 5 uniquely upregulated and downregulated proteins in assembloids and mosaics are shown (Figure 5B). GO enrichment analyses revealed the association of pathway regulation with each co-culture condition (Figure 5C). Interestingly, the only biological process identified as being significantly upregulated in DIPG cells from mosaics was the GO term nBAF, a process linked to the regulation of genes that are essential for dendrite growth. Collectively, the differential pathway activation in DIPG cells grown as mosaics versus assembloids highlights potential pathways that are upregulated when DIPG cells interact directly with cells of the cortical organoid.

**Figure 5.**
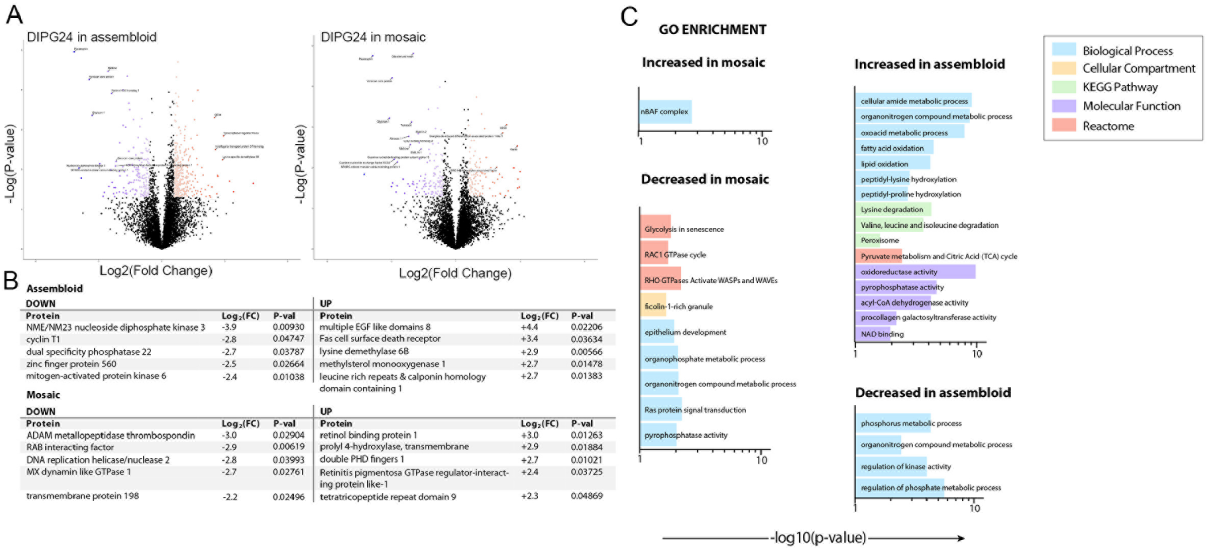
Analysis of mosaic cultures reveals unique signalling in DIPG cells in contact with normal brain cells. A. Volcano plots showing expression changes (log2[foldchange]) in DIPG24 cells in assembloid and mosaic cultures compared to DIPG24 cells grown as spheroids and respective-log(p-value) for each fold change. Significant changes (*P*<0.05) are highlighted in colour where increases are shown in orange and decreases in blue. Colour gradient correlates with the degree of fold change, with the greater change shown in darker colours. B. Table shows the top 5 altered genes (either increased or decreased expression) for each condition shown in (A). C. Gene ontology (GO) enrichment analyses for the indicated expression changes. Bar charts show significance (-log10[p-value]) of each term.

## DISCUSSION

Discovery of successful therapies for DIPG requires models that mimic *in vivo* intra-cellular interactions and support invasive dissemination. In the present study we demonstrate that cortical organoids provide a permissive environment for DIPG invasion and that co-culture triggers key signalling changes. Analysis of changes specific to the dispersed DIPG cells from mosaics unveiled unique activation of dendritic growth pathways. The demonstration of invasive phenotypes and proteome responses suggest that DIPG brain organoid co-cultures are a valuable tool for analysing DIPG biology that may be exploited for treatment.

A major goal of our study was to establish a model that replicated DIPG invasion and dissemination. DIPG cells indeed dispersed throughout the organoids, occasionally incorporating into neural rosettes, confirming the intimate interactions between the tumour and organoid cells. It has previously been shown that GBM similarly avidly invade cerebral organoids in co-culture ^17, 20^. Brain organoids therefore provide a species-specific *in vitro* model for investigating the dissemination of high-grade gliomas including GBM and DIPG. The present study used spheroids prepared from DIPG cells grown under defined media conditions lacking serum and kept at low passage to maintain stem cell populations and genomic integrity. In the progress towards ever-more-faithful cell cultures glioblastoma organoids (GBOs), where small pieces of resected tumour are cultured *ex vivo* ^21, 22^, have recently been developed. GBOs maintain genome integrity, retain important inter- and intra-tumoural heterogeneity and for a short period of time retain the mix of normal cell types characteristic of the *in vivo* tumour microenvironment. However, extending GBO models to the investigation of invasion requires orthotopic xenografts ^21^, a more costly and non-human tissue experimental model. In future, combination of GBOs (or DIPG organoids) with cortical organoids, could enhance the *ex vivo* tumour organoid approach to facilitate analysis of invasion. The organoid/DIPG co-culture method is cheaper and faster than mouse models, provides a species-relevant tissue setting and 3D context that is lacking in 2D models. Specific questions concerning regional tropism could be further interrogated by introducing spheroids/GBOs to region-specific brain organoids ^23, 24^.

SWATH-MS proteomic profiling of tumour cell populations revealed a marked enrichment of DNA replication and mitotic activity GO terms in tumour cells cultured in organoid co-culture. This highlights the importance of the healthy brain microenvironment in tumour progression and reinforces the rationale for cortical organoids as a pre-clinical model. Limited available DIPG tissue has led to a focus on gene expression rather than proteomic analyses ^25, 26^. Importantly, it is easy to differentially label the brain organoid and tumour cells, facilitating separation of the different cellular populations for proteomic analysis. Given the frequent lack of correlation between gene transcripts and eventual protein levels that results from the extensive array of post-transcriptional mechanisms, capturing the proteomic profiles is an important addition to the signalling landscape of DIPG.

Pathway analyses suggested that DIPG adhesion was decreased in co-culture and this was confirmed by immunostaining. The altered distribution of integrin suggested a decrease in adhesive contacts between DIPG cells. This has parallels in the altered morphology and adhesion that accompanies progression to invasive carcinoma ^27^.

Previous studies have shown that GBM/brain organoid mosaics stimulates gene profiles associated with network formation and invasion ^17^. Data from the present study comparing the assembloid and mosaic models revealed that significant changes occur simply due to co-culture with organoids, without requirement for direct contact between the DIPG and organoid cells. By filtering out the proteomic changes common to both assembloids and mosaics we pinpointed the changes that were instigated by contact between DIPG and organoid cells. The data suggested that contact with organoid cells upregulated neuron-specific brahma-associated factor (nBAF) complex signalling. The nBAF complex is a developmentally regulated complex during normal development, precisely timed to function during the maturation of neuronal progenitors as they exit the subventricular zone of the brain ^28^. Contact between DIPG and cortical organoids cells may upregulate this developmentary regulated pathway that is associated with neuronal cell invasion.

Tumour biology is determined by interactions between tumour cells, between tumour cells and normal cells, and between tumour cells and surrounding extracellular components ^13^. This is particularly relevant for DIPG where disease progression is closely tied to tumour-normal cell interactions. The extent of diffuse infiltration into healthy brain tissue is a key indicator of worsening prognosis^1^. *In vitro* 2D models do not account for tumour-tumour and -normal cell interactions, nor the role of extracellular components. Genetic mouse and PDX models offer a solution but raise the issue of interspecies differences ^29^, along with the ethical imperative to reduce use of animal models where appropriate. Here we have described a DIPG-cortical organoid co-culture which overcomes many of these limitations. Most importantly, the organoids provide a permissive environment for DIPG invasion and dissemination. Since invasion and dissemination are a major impediment to successful DIPG treatment, organoid co-culture offers the opportunity to better understand this pathological phenotype and thereby derive improved therapies.

## Materials and methods

### Cell culture

Primary patient-derived DIPG cell line, DIPG24, was kindly supplied by Michelle Monje (Stanford University, USA ^5^). Where indicated, DIPG24 cells transfected with lentivirus encoding GFP and selected using puromycin were used. Tumour cells were cultured as an adherent monolayer in Tumour Stem Media (TSM) containing Neurobasal^TM^ without vitamin A Medium, DMEM F-12, 10 mM HEPES, 1x B27^TM^ without vitamin A supplement, 1 mM Sodium pyruvate, 1x MEM Non-essential amino acids solution, 1x GlutaMAX^TM^, 1x Antibiotic-Antimycotic (all Life Technologies, MA, USA), supplemented with 20 ng/mL EGF and FGF2, 10 ng/mL each of PDGF-AA and PDGF-BB (Shenandoah Biotechnologies, PA, USA). All cultures were maintained at 37°C 5% CO2. H9 human embryonic stem cells (Agreement No. 19-W0538; WiCell, USA) were maintained as bulk culture in feeder-free conditions on vitronectin (StemCell Technologies, Canada)-coated dish in TeSR-E8 basal medium plus supplement (StemCell Technologies, Canada). Stem cells were karyotyped to confirm genomic stability (Sullivan Nicoloaides Pathology, Brisbane, Australia). No abnormalities were detected in Geimsa stained metaphase spreads from 15 cells, at 300 bands per haploid set. All stem cell culture and procedures were in accordance with guidelines of the Sydney Children’s Hospital Network (2019/ETH00240) and University of Wollongong Health and Medical Human Research Ethics Committee (2020/451). All cell lines were short tandem repeat (STR) profiled to confirm identity (CellBank Australia) and limited to a maximum of 20 passages after STR profiling.

### Dorso-cortical organoid formation

H9 stem cells (WiCell, WI, USA) and ENVY HES3 stem cells^30^ (kindly provided by Andrew Elefanty, Murdoch Children’s Research Institute, VIC, Australia) were passaged using 0.5 mM EDTA (Life Technologies, MA, USA), and plated on laminin (Life Technologies, MA, USA)-coated plates in TeSR-E8 basal medium. After 24 hours, medium was replaced with Neural induction medium (NIM) (Neurobasal^TM^ Medium, DMEM F-12, 1x N2 supplement, 1x B27^TM^ without vitamin A supplement, 1x ITS without vitamin A, 2 mM GlutaMAX ^TM^, 0.3% (w/v) glucose) (Life Technologies, MA, USA)), supplemented with SMAD inhibitors 0.1mM LDN193189 (StemCell Technologies, Canada), 10μM SB431524 (StemCell Tecnologies, Canada). Media was replenished every one-two days over a period of six days, where neural rosettes formed on top of colonies. On day six, loosely adhered rosettes were dissociated using 0.5 mM EDTA, harvested, and plated as aggregates in an ultra-low attachment (ULA) U-bottom 96-well plate (Corning, USA) in ‘Neural Media’ (NM) (Neurobasal ^TM^ Medium, 1x N2 Supplement, 1x B27 ^TM^ without vitamin A supplement, 1x ITS without vitamin A, 2 mM GlutaMAX ^TM^ (Life Technologies, MA, USA)) supplemented with 20 ng/mL EGF and 20 ng/mL FGF2 (Shenandoah Biotechnologies, PA, USA) (growth factor-supplemented NM). Neurospheres were cultured under static conditions for 14 days before moving to a ULA 6-well plate (Corning, USA) and cultured in an orbital shaker incubator at 85-100 RPM with a 2.5 cm orbital diameter to prevent necrosis in the centre of the cultures. Neurospheres matured to organoids over 10-14 days, with NM supplemented with EGF and FGF2 changed every 2-3 days. GFP-positive ENVY cells were used for organoid generation in proteomic analyses and initial tests of co-culture conditions (eg Supplementary Figure 1). Otherwise, H9-derived organoids were used to validate findings and in the analysis of organoid and co-culture structures.

### Tumour spheroid formation

Cells were detached from their tissue culture flasks with Accutase^TM^ solution (Corning, USA), and plated at a density of 5,000 cells in 100 µL cell culture medium per well of a non-adhesive, U-bottom, 96-well dish (Corning, USA). Spheroids formed and compacted over 5 days during incubation at 5% CO2 37°C. Where indicated, cells were first transduced with a second-generation lentivirus expressing both eGFP and luciferase.

### Dorsal-cortical organoid co-culture with tumour spheroids

At 16 to 20-days post-induction, when organoids exhibited typical features including formation of neural rosettes and expression of dorsal cortical markers identified by immunohistochemistry including Vimentin, SOX2 and PAX6, TBR2, βTUBIII, and MAP2 ^31–36^, organoids were harvested for co-culture. Tumour spheroids were collected 5 days post-seeding. Organoids and tumour spheroids were collected separately and rinsed in 1x DPBS (Life Technologies, MA, USA). For assembloid cultures, organoids and tumour spheroids were plated in an ULA U-bottom 96-well plate at a 1:1 ratio in growth-factor supplemented NM. For mosaic cultures, individual organoids and tumour spheroids were dissociated by incubation with Accutase^TM^ 37°C for 10 minutes, followed by the addition of an equal volume of trypsin inhibitor. Cells were then pelleted by centrifugation at 200xg for three minutes and resuspended in 50 μL growth factor supplemented NM per spheroid or organoid. 50 μL of each cell suspension was then combined at a 1:1 ratio in a ULA U-bottom 96-well plate. Immediately following plating, assembloid and mosaic culture plates were centrifuged at 200 xg for 3 minutes, and returned to incubation at 5% CO2, 37°C. After 3 days, co-cultures were transferred to ULA 6-well plate (10 co-cultures per well) in an orbital shaking incubator at a speed of 85-100 RPM with an orbital diameter of 2.5 cm. For proteomic analyses, all cultures, including tumour spheroid controls, were grown in NM.

### Immunofluorescence

Cultures were rinsed in DPBS and fixed with 4% paraformaldehyde (PFA) solution at 4°C, for 1 hour and further incubated in 20% (v/v) sucrose solution overnight at 4°C. Samples were mounted using Optimal cutting temperature (OCT) compound (Sakura Finetek, CA, USA), and frozen at −80°C until sectioning. OCT-embedded samples were cryosectioned to a thickness of 15 μm and stored frozen at −20°C until processed for immunofluorescence. Slide-mounted sections were thawed and rinsed in DPBS to remove OCT. Regions of interest for staining were defined using a hydrophobic PAP pen (Sigma, USA). Sections were permeabilized with 0.02% (v/v) Triton X-100/PBS for 10 minutes before blocking for 1 hour with blocking buffer (10% (v/v) donkey serum/PBS). Primary antibodies for staining included vimentin (1/100 dilution, Santa Cruz Biotechnology, TX, USA), SOX2 (1/200 dilution, Cell Signalling, MA, USA), PAX6 (Developmental Studies Hybridoma Bank, IA, USA), TBR2 (1/50 dilution, Novus Biologicals, CO, USA), βTUBIII (1/200 dilution, Cell Signalling Massuchusetss, USA), MAP2 (1/400 dilution, Sigma-Aldrich, Germany), Ki-67 AlexaFluor 647 conjugated (1/100 dilution, BD Biosciences, NJ, USA), cleaved Caspase 3 (1/200 dilution, Cell Signalling, MA, USA) and β1 integrin (1/50 dilution; Novus Biologicals, CO, USA). Primary antibodies diluted in blocking buffer were incubated overnight at 4°C. After rinsing with 0.05% Triton-X 100/PBS (PBS-T), sections were incubated with fluorophore-conjugated secondary antibodies (Invitrogen, MA, USA) diluted in 10% (v/v) donkey serum/PBS for one hour at 4°C. Sections were rinsed and incubated with blocking buffer containing 10 μg/mL Hoechst blue (Life Technologies, MA, USA) for 20 minutes. Sections were rinsed with PBS-T and once with deionized water before mounting with FluorSave mounting medium (Merck-Milipore, Germany) and coverslipping. Cells were imaged using a Leica TCS SP5 inverted confocal microscope (Leica Biosystems, Germany).

### Western blotting

Cells, organoids or tumour spheroids were lysed on ice in a Sodium dodecyl sulfate (SDS)-Radioimmunoprecipitation (RIPA) buffer (50 mM Tris pH 7.4, 150 mM NaCl, 5 mM EDTA pH 8.0, 1% (v/v) Nonidet P-40, 0.1% (w/v) SDS, 1% (w/v) sodium deoxycholate) supplemented with protease inhibitors (1mM Na3VO4, 1 mM phenylmethanesulfonyl fluoride, 1 μg/mL aprotinin, 1 μg/mL leupeptin). Samples were pulse sonicated, and then centrifuged at 15 500xg to remove cellular debris and non-solubilised proteins. 5 to 30 μg of each lysate was eletrophoretically separated in a polyacrylamide gel and electroblotted onto a Polyvinylidene difluoride (PVDF) membrane. Membranes were incubated with primary antibodies including OCT4 (1/2000 dilution, Abcam, MA, USA), and SOX2 (1/1000 dilution), BTUBIII (1/500 dilution) and GAPDH (1/5000 dilution, Invitrogen, MA, USA). Immunoreactive proteins were detected by chemiluminescence using horseradish peroxidase-conjugated secondary antibodies (Invitrogen, MA, USA) and ECL plus reagent (Merck Millipore, MA, USA), detected using an Odessey Fc imaging system (LICOR, USA).

### Cell sorting

Co-cultured cells were collected, rinsed in DPBS, and dissociated by gentle trituration following incubation in Accutase^TM^ at 37°C for 10 minutes. Cells were resuspended in cold FACS buffer (137.4 mM NaCl, 2.7 mM KCl, 10.1 mM Na_2_HPO_4_, 1.8mM KH2PO4, 0.1% (w/v) bovine serum albumin (BSA)) and kept on ice. Cell suspensions were passed through a 35 μm PES cell strainer (Corning, USA) and spiked with 1 μg/mL DAPI (Sigma Aldrich, Germany) directly prior to analysis on a FACSAria II (BD Biosciences, USA). Spectral data was processed using the FACSDiva Software Package (BD Biosciences, NJ, USA). Debris, doublets, and dead cells were excluded by manual gating. GFP+ve and GFP-ve populations were sorted and collected in cold NM. Cells were pelleted by centrifugation at 200 xg for 3 minutes, snap frozen on liquid nitrogen, and stored in liquid nitrogen for later protein extraction and digestion.

### SWATH Proteomics

Label-free, next-generation proteomic Sequential Window Acquisition of all Theoretical Mass Spectra (SWATH-MS) was used to resolve proteomes of tumour and organoid populations (Ludwig et al. 2018). Snap-frozen cell pellets were resuspended in extraction buffer (1% (w/v) SDS, 8 M urea, 100 mM Tris-HCl (pH 8.0)) and sonicated using a Sonifer250 (Branson, USA) to extract proteins, followed by centrifugation to remove debris. The protein concentration for each sample was determined by BCA protein assay kit (Thermo Fisher Scientific, USA). Cysteine disulphide bonds in the proteins were reduced with 10 mM dithiothreitol (DTT) at 37°C for one hour, and then alkylated with 20 mM iodoacetamide (IAA) for 45 minutes in the dark at room temperature. IAA was then quenched with 10 mM DTT for 15 minutes in the dark at room temperature. Proteins were digested with trypsin using S-TrapTM Micro spin columns, as per manufacturer’s instructions. Briefly, an equal volume of 2x lysis buffer (10% (w/v) SDS, 50 mM Triethylammonium bicarbonate (TEAB) pH 7.55) was added, followed by sample acidification with aqueous H3PO4 at a final concentration of ∼1.2% (v/v). S-Trap^TM^ binding buffer (90% (v/v) methanol, 100 mM TEAB, pH 7.1) was then added to the acidified sample (6:1, binding buffer:sample), and this mixture was passed through the micro spin column followed by three washes with binding buffer. Trypsin proteo (10:1, sample:trypsin) was performed on S-Trap^TM^ columns at 47 °C for one hour. Digested peptides were eluted with 50 mM TEAB, followed by 0.2% (v/v) formic acid, and lastly with 50% (v/v) acetonitrile in 0.2% (v/v) formic acid. Eluted peptides were vacuum dried and resuspended in loading buffer (2% (v/v) acetonitrile 0.1% (v/v) formic acid). The peptide concentration for each sample was determined by peptide BCA assay.

### Liquid Chromatography & Mass Spectroscopy

Two pools were prepared from each organoid and tumour sample (total of 120 μg), respectively. These were fractionated by high pH reverse phase high performance liquid chromatography (RP-HPLC) using an Eksigent Ultra nanoLC system (Sciex, USA). The dried digested sample was resuspended in mobile phase buffer A (5 mM ammonium hydroxide solution, pH 10.5). After sample loading and washing with 3% (v/v) buffer B (5 mM ammonia solution with 90% acetonitrile, pH 10.5) for 10 minutes at a flow rate of 300 μL/minute, the buffer B concentration was increased from 3% to 30% over 55 minutes and then to 70% between 55 to 65 minutes and to 90% between 65-70 minutes. The eluent was collected every 2 minutes at the beginning of the gradient and at one-minute intervals for the rest of the gradient. Following high pH-RP-HPLC separation, 17 fractions were concatenated (0–82 minutes), dried and resuspended in 20 μL of loading buffer. 10 μL/fraction was taken for two-dimensional information dependent acquisition (IDA) analysis. 2 μg of the cleaned sample was taken and diluted with the loading buffer to a final volume of 10 μL for SWATH analysis. SWATH were acquired in random with blank runs in between samples.

### Proteomic data Acquisition

Sample (10 μL) was injected onto a reverse-phase C18 self-packed peptide trap (Halo-C18, 160 Å, 2.7 μm, 200 μm x 10 mm) for pre-concentration and desalted with loading buffer, at 5μL/minute for 3 minutes (Triple TOF 6600; Sciex, USA). The peptide trap was then switched into line with the analytical column (15 cm x 200 μm nano cHiPLC column (ChromXP C18-CL 3μm 120 Å)). Peptides were eluted from the column using a linear solvent gradient from mobile phase A: mobile phase B (95:5) to mobile phase A: mobile phase B (65:35) at 600 nL/minute over a 120-minute period. After peptide elution, the column was cleaned with 95% buffer B for six minutes and then equilibrated with 95% buffer A for 10 minutes before next sample injection. The reverse phase nanoLC eluent was subject to positive ion nanoflow electrospray analysis in an IDA mode. In the IDA mode a time-of-flight mass spectrometry (TOF-MS) survey scan was acquired (m/z 350-1500, 0.25 second) with the 20 most intense multiply charged ions (2+-5+; exceeding 200 counts per second) in the survey scan sequentially subjected to MS/MS analysis. MS/MS spectra were accumulated for 100 milliseconds in the mass range m/z 100–1800 with rolling collision energy optimized for lowed m/z in m/z window +10%.

### Proteomic data processing and analysis

Raw data files generated by IDA-MS analysis were searched with ver5.0 ProteinPilot (Sciex, USA) using the Paragon TM algorithm in thorough mode. Homo sapiens species from SwissProt (SwissProt_2019_05.fasta) containing 20, 420 proteins was used for searching the data. Carbamidomethylation of cysteine residues was selected as a fixed modification, and digestion with trypsin was used. An Unused Score cut-off was set to 1.3 (95% confidence for identification), and global protein false discovery rate (FDR) of 1%. A spectral ion library was constructed by merging all the 2D-IDA libraries from both tumour and organoid samples. SWATH data were extracted using ver 2.2 PeakView (Sciex, USA). The top 6 most intense fragments of each peptide were extracted from the SWATH data sets (75 parts per million (ppm) mass tolerance, 10-minute retention time window). Shared and modified peptides were excluded. After data processing, peptides (max 100 peptides per protein) with confidence ≥99% and FDR ≤1% (based on chromatographic feature after fragment extraction) were used for quantitation. The extracted SWATH protein peak areas were analysed using an in-house software program (APAF, Macquaire University). Protein peaks were normalised to the total peak area for each run and subjected to two sample *t*-test to compare relative protein peak area between the sample groups. Protein peaks with value < 0.05 and fold change ± 1.5 were deemed as differentially expressed proteins. The fold change in protein abundance was compared between different culture conditions. Protein interaction networks were identified using the STRING database (https://string-db.org) ^37^ and gene ontology term enrichment analyses performed using G-profiler (https://biit.cs.ut.ee/gprofiler/gost) ^38^.

### RT-PCR

Custom primers for real-time PCR were purchased from Sigma-Aldrich, Germany (GAPDH: forward 5’TGCACCACCAACTGCTTAGC3’, reverse 5’GGAAGGCCATGCCAGTGA3’; MCM2: forward 5’CACAACGTCTTCAAGGAGCG3’, reverse 5’TCGTACTTGGGGTACATGGC3’; NDC1: forward 5’ATTCCCAAAGCTTGGATTAGCA3’, reverse 5’GACATACCAAGTCGTCAGGAG3’; NUP43: forward 5’TGACCAGGAAAGAATTGTCGC3’, reverse 5’GGTGCACTGCTATAGGAAGGA3’; NUP50: forward 5’GAAGGTGACAGTGGTGAATG3’, reverse 5’AAATGCAGAGTACCTATGCC3’; NUP107: forward 5’GGCTGGAAACTGTACCATGAC3’, reverse 5’GGAAGTAGGCCCAAACTGTG3’; POLA2: forward 5’TGGCAGCGAACTCAAGGAAC3’, reverse 5’CGAGGAATGTTCCCGGTCTC3’). Total RNA was extracted with TRIzol^TM^ LS reagent as per manufacturer’s instruction, quantified using a NanoDrop ^TM^ 2000 spectrophotometer (Thermo Scientific, USA), and reverse transcribed with the SuperScript^®^ III first-strand synthesis system (Invitrogen, MA, USA). Quantitative PCR was performed with iTaq universal SYBR green supermix (BioRad, CA, USA) using the CFX96 Real Time system (BioRad, CA, USA) in the default thermal cycling mode. GAPDH was used as a normalisation reference. All reagents were used according to the manufacturer’s instructions. The 2-ΔΔCT method was used as a relative quantification strategy for quantitative real-time PCR data analysis. RT-PCR analyses were performed using CFX Maestro software ver1.1 (Bio-Rad, CA, USA).

## ACKNOWLEDGEMENTS

Aspects of this research have been facilitated by access to the Australian Proteome Analysis Facility (APAF, Macquarie University, AUS) supported under the Australian Government’s National Collaborative Research Infrastructure Strategy (NCRIS). We would like to thank APAF for their assistance with SWATH MS and proteomic analyses, and Westmead Hub Flow Facility for their assistance with cell sorting. We acknowledge Dr Anai Cordero Gonzalez (Children’s Medical Research Institute, AUS) for helpful advice. We thank Daniel Wallis for his assistance with generating Python scripts for *in silico* data analyses. We gratefully acknowledge funding support from The Kids Cancer Project/Perpetual IMPACT Funding, the Balance Foundation, and a Mid-Career Research Accelerator grant from the University of Sydney to GON. VP was supported by an Australian Government Research Training Program (RTP) Scholarship and a Top-up Scholarship from The Kids Cancer Alliance.

## COMPETING INTERESTS

The authors declare no competing financial interests

## Notes

### Competing Interest Statement

The authors have declared no competing interest.

