## Supplementary Figs 1 - 3 for "Parsing the effect of co-culture with brain organoids on Diffuse Intrinsic Pontine Glioma (DIPG) using quantitative proteomics"

Supplementary Figure 1

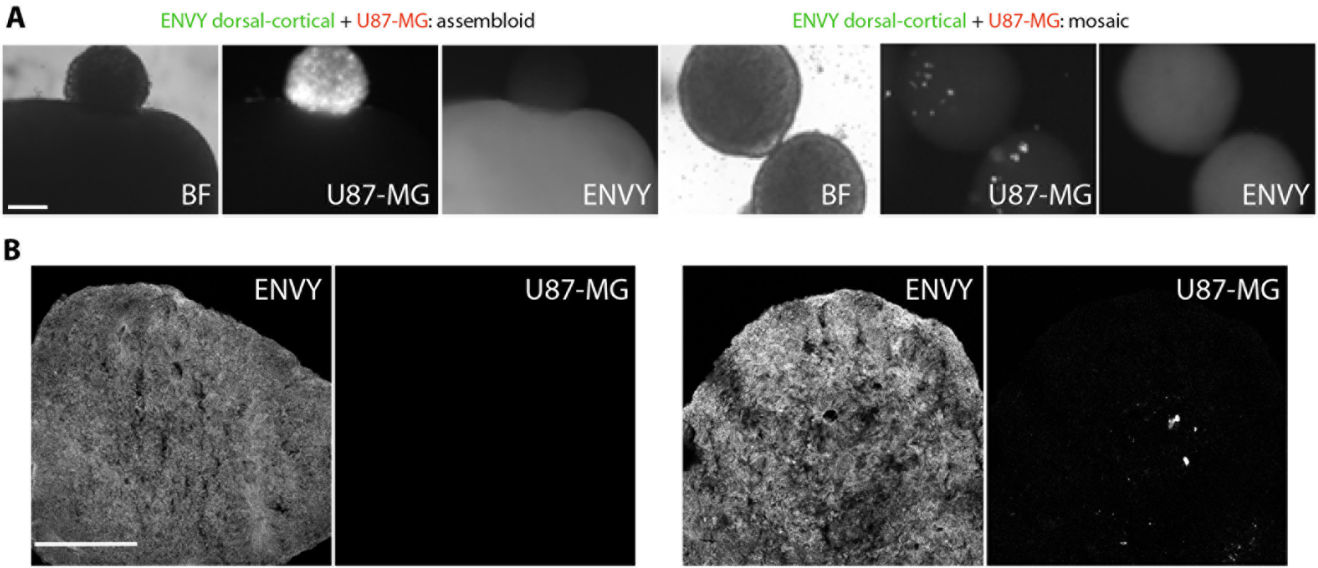


**Tumour spheroids of U87-MG do not form assembloid or mosaic co-cultures with Envy dorsal cortical organoids.** Dorsal cortical oganoids induced from GFP-expressing hESC line Envy were cultures with U87-MG spheroids labelled with Cell Tracker Orange. U87-MG spheroids did not form assembloid cultures and tumour cells did not invade organoids. **A**. Time-lapse brightfield (BF) and epifluorescence imaging shows U87-MG spheroid in contact with an organoid. **B**. Maximum intensity z-projections of confocal images of sections organoids confirms the absence of any U-87MG cells in the assembloid organoid and miminmal incorporation of U87-GM tumour cells in mosaic organoids. Scale = 200 µm. Images shown are representative examples from 3 independent replicates

### Supplementary Figure 2

#### A Assembloid co-cultures

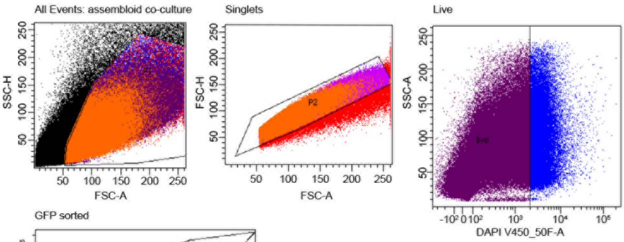


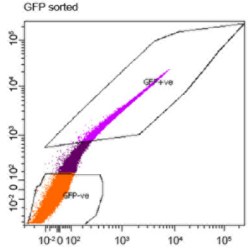


| Population | #Events | %Parent | %Total |
|---|---|---|---|
| All Events | 1,000,000 | ### | 100.0 |
| P1 | 418,174 | 41.8 | 41.8 |
| P2 (singlets) | 393,811 | 94.2 | 39.4 |
| Live cells | 348,966 | 88.6 | 34.9 |
| GFP+ve (organoid) | 102,573 | 29.4 | 10.3 |
| GFP-ve (tumour) | 211,266 | 60.5 | 21.1 |

#### Mosaic co-cultures

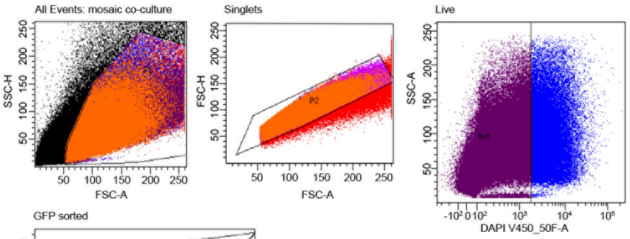


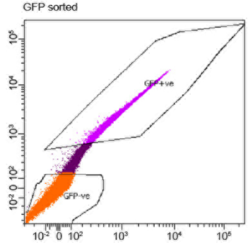


| Population | #Events | %Parent | %Total |
|---|---|---|---|
| All Events | 1,000,000 | ### | 100.0 |
| P1 | 534,243 | 53.4 | 53.4 |
| P2 (singlets) | 501,311 | 93.8 | 50.1 |
| Live cells | 445,726 | 88.9 | 44.6 |
| GFP+ve | 132,098 | 29.6 | 13.2 |
| GFP-ve | 307,057 | 68.9 | 30.7 |

## B

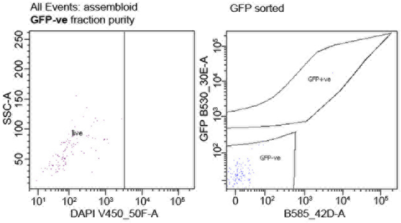


| Population | %Parent | %Total |
|---|---|---|
| All Events | ### | 100.0 |
| P1 | 22.7 | 22.7 |
| P2 (singlets) | 99.2 | 22.5 |
| Live cells | 100.0 | 22.5 |
| GFP+ve | 1.0 | 0.2 |
| GFP-ve | 99.0 | 22.2 |

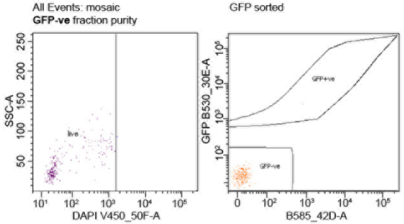


| Population | %Parent | %Total |
|---|---|---|
| All Events | ### | 100.0 |
| P1 | 43.6 | 43.6 |
| P2 (singlets) | 99.6 | 43.5 |
| Live cells | 99.6 | 43.3 |
| GFP+ve | 0.4 | 0.2 |
| GFP-ve | 99.6 | 43.1 |

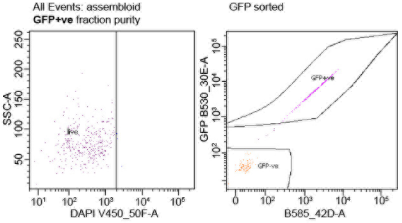


| Population | %Parent | %Total |
|---|---|---|
| All Events | ### | 100.0 |
| P1 | 77.7 | 77.7 |
| P2 (singlets) | 98.9 | 76.8 |
| Live cells | 99.1 | 76.2 |
| GFP+ve | 70.4 | 53.6 |
| GFP-ve | 27.2 | 30.8 |

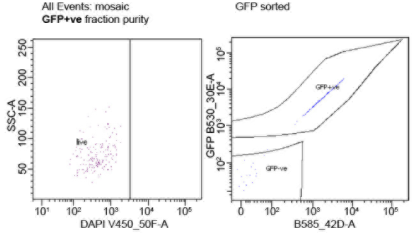


| Population | %Parent | %Total |
|---|---|---|
| All Events | ### | 100.0 |
| P1 | 57.7 | 57.7 |
| P2 (singlets) | 98.8 | 57.0 |
| Live cells | 100.0 | 57.0 |
| GFP+ve | 89.0 | 50.7 |
| GFP-ve | 10.4 | 5.9 |

**Fluorescence activated cell sorting separates co-cultures into individual tumour and organoid cell fractions. A** Dissaggregated assembloid and mosaic co-cultures were FACS-sorted on the basis of GFP expression. Plots show cells sorted using a hierarchical gating strategy, where a single cell population was assesed for viability using DAPI staining. Live-single cells were then gated on the basis of GFP expression to yield separate GFP+ve organoid, and GFP-ve tumour cell populations. Colours of events in plots refer to populations represented in table, where indents of populations represent hierarchical 'daughter' gates. **B** Purity of GFP+ve (organoid) and GFP-ve (tumour) fractions was tested by running sorted samples through the same hierarchial gating used during sorting (only live and GFP-gated plots shown).

### Supplementary Figure 3

**A**

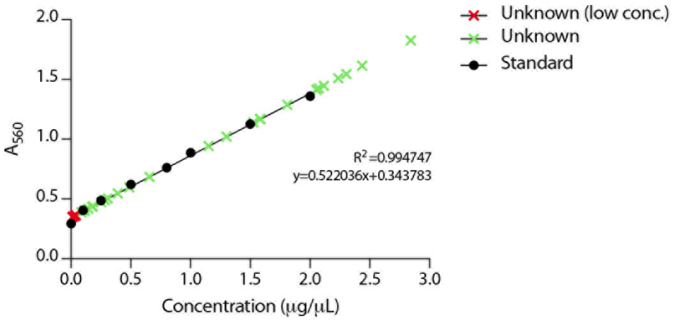


| | Replicate | | | |
|---|---|---|---|---|
| **Condition** | **1** | **2** | **3** | **4** |
| Assembloid-control organoid | × | × | × | × |
| Mosaic-control organoid | × | × | × | × |
| Spheroid | × | × | × | × |
| Spheroid (TSM) | × | × | × | × |
| Assembloid GFP+ve organoid | × | × (red) | × (red) | × (red) |
| Assembloid GFP-ve tumour | × | × | × | × (red) |
| Mosaic GFP+ve organoid | × (red) | × (red) | × | × (red) |
| Mosaic GFP-ve tumour | × | × | × | × |

**B**

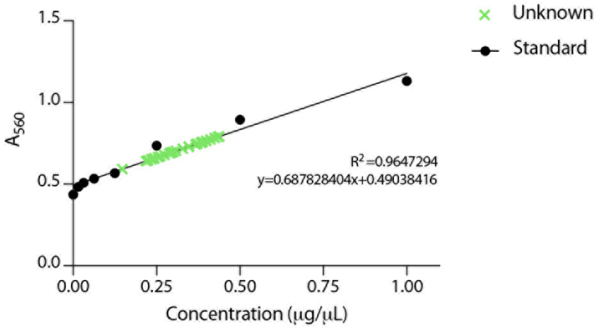


**C**

| Sample | Library Bin |
|---|---|
| Spheroid Replicate 1 | Tumour |
| Spheroid Replicate 2 | Tumour |
| Spheroid Replicate 3 | Tumour |
| Spheroid Replicate 4 | Tumour |
| Spheroid TSM Replicate 1 | Tumour |
| Spheroid TSM Replicate 2 | Tumour |
| Spheroid TSM Replicate 3 | Tumour |
| Spheroid TSM Replicate 4 | Tumour |
| Assembloid GFP-ve tumour Replicate 1 | Tumour |
| Assembloid GFP-ve tumour Replicate 2 | Tumour |
| Assembloid GFP-ve tumour Replicate 3 | Tumour |
| Assembloid GFP-ve tumour Replicate 4 | Tumour |
| Mosaic GFP-ve tumour Replicate 1 | Tumour |
| Mosaic GFP-ve tumour Replicate 2 | Tumour |
| Mosaic GFP-ve tumour Replicate 3 | Tumour |
| Mosaic GFP-ve tumour Replicate 4 | Tumour |
| Assembloid-control organoid Replicate 1 | Organoid |
| Assembloid-control organoid Replicate 2 | Organoid |
| Assembloid-control organoid Replicate 3 | Organoid |
| Mosaic-control organoid Replicate 1 | Organoid |
| Mosaic-control organoid Replicate 2 | Organoid |
| Mosaic-control organoid Replicate 3 | Organoid |
| Mosaic-control organoid Replicate 4 | Organoid |
| Assembloid GFP+ve organoid Replicates pooled | Organoid |
| Mosaic GFP+ve organoid Replicates pooled | Organoid |

**Protein quantification for digestion and concentration for high-pressure liquid chromatography and peptide library assembly.** Protein from disaggregated, fluorescence activated cell sorted (FACS)-cultures was extracted with a 1% (w/v) SDS, 8M Urea, 100mM Tris HCl solution, and purified following a chloroform/methanol wash. **A** Resulting protein was quantified by BCA assay by measuring the absorbance of unknowns against a standard curve of known protein concentrations, generated from a linear regression fit of standard samples. Graph shows $R^2$ and equation of the curve. Samples with total amount of protein of less than 20µg (red cross) were pooled into a single sample for each indicated condition for further protein digestion and concentration. **B** Resulting peptides were quantified by BCA assay against a standard curve of known peptide concentrations, as described above. **C** Equal amounts of digested protein from samples were pooled into a tumour or organoid library according to cell type of origin.
